# Decoding the dynamics of cognitive control: Insights from reach movements and electroencephalography

**DOI:** 10.1101/2025.08.29.673147

**Authors:** Moaz Shoura, Katie McNair, Adrian Nestor, Christopher Erb

**Author notes:** Correspondence concerning this article should be addressed to Moaz Shoura, Department of Psychology at Scarborough, University of Toronto, 1265 Military Trail, Scarborough, Ontario, Canada, M1C 1A4. These authors contributed equally (Moaz Shoura, Katie McNair). Present affiliation: Department of Neurology, Boston Children’s Hospital (Katie McNair). Declarations Funding. Christopher Erb and Adrian Nestor disclose support for this research from the Marsden Fund, administered by the Royal Society Te Apārangi. Adrian Nestor discloses support from the Natural Sciences and Engineering Research Council of Canada (NSERC). **Conflict of Interest Statement.** The authors declare no competing interests. Ethics approval. All procedures were approved by Human Participants Ethics Committee (UAHPEC) at the University of Auckland. **Consent to participate.** Each participant provided informed consent and received monetary compensation for their involvement. **Consent for publication.** N/A. **Availability of data and materials.** All data and stimuli have been made publicly available via the Open Science Framework and can be accessed at: https://osf.io/cdj2z/. None of the reported studies were preregistered. **Code availability:** All code has been made publicly available via the Open Science Framework and can be accessed at: https://osf.io/cdj2z/. **Authors’ contributions. Moaz Shoura:** Conceptualization, Methodology, Software, Formal analysis, Validation, Data Curation, Writing – Original Draft, Writing – Review & Editing, Visualization. **Katien McNair:** Conceptualization, Methodology, Investigation, Formal analysis, Data Curation, Writing – Review & Editing. **Adrian Nestor:** Conceptualization, Methodology, Writing – Review & Editing, Supervision, Project administration, Funding acquisition. **Christopher Erb:** Conceptualization, Methodology, Investigation, Writing – Review & Editing, Resources, Supervision, Project administration, Funding acquisition.

## Abstract

The congruency sequence effect (CSE) refers to improved performance in conflict tasks, such as the Eriksen flanker task, when the congruency of the current trial matches that of the preceding one. Although widely attributed to dynamic adjustments in cognitive control, the neural mechanisms and temporal dynamics underlying this effect remain poorly understood. Here, we combined a release-and-press flanker task with EEG decoding to examine how trial congruency and response type shape behavior and brain activity in healthy adults. Behaviorally, the CSE emerged only when responses repeated, highlighting the dominant role of stimulus-response binding over abstract control mechanisms. Neural decoding mirrored the CSE: current-trial congruency was more reliably decoded on repeated-response trials following congruent versus incongruent trials. This neural CSE was stimulus-locked, occurred between 450-550ms post-stimulus onset, and was driven by theta-band activity. More broadly, congruency decoding was particularly robust over frontal channels while cross-temporal generalization indicated transient, sequential neural representations underlying control signals. Together, these findings demonstrate that both behavioral and neural signatures of the CSE are tightly constrained by response repetition and emerge within a narrow temporal window. By integrating reach-based measures with multivariate EEG analyses, this study provides a fine-grained account of when, where, and under what conditions control-related signals unfold after conflict.

## Introduction

Cognitive control enables flexible, goal-directed behavior by aligning actions and thoughts with goals and contextual demands (Badre, 2008; Miller & Cohen, 2001; Shenhav et al., 2013). Classic paradigms that require overriding habitual responses amidst conflicting cues, such as the Stroop (Stroop, 1935), Simon (Simon, 1969) and flanker (Eriksen & Eriksen, 1974) tasks, have been central to investigating the cognitive and neural mechanisms of control (Botvinick et al., 2001). In these tasks, a congruency effect is reflected in slower and more error-prone responses on incongruent trials (Kornblum et al., 1990; Ridderinkhof et al., 1995).

Further complexity is revealed through congruency sequence effects (CSEs), whereby congruency effects are reduced following incongruent compared to congruent trials (Egner, 2007; Gratton et al., 1992). Initially interpreted as evidence of conflict-driven adaptive control (Botvinick et al., 2001, 2004; Kerns et al., 2004), the CSE has also been attributed to associative priming and stimulus–response (S–R) binding, where perceptual and motor features are temporarily bound into an “event file” and automatically retrieved when any of those features recur (Hommel, 2004; 2005). However, while controlling for stimulus repetitions can reduce the CSE (Nieuwenhuis et al., 2006; Schmidt & De Houwer, 2011), effects may persist even when feature repetitions and contingency biases are eliminated (Kim & Cho, 2014; Schmidt & Weissman, 2014). Consequently, contemporary models emphasize a dynamic interplay between memory-driven bindings, including S–R integration, and genuine top-down control (Dignath et al., 2019; Egner, 2017).

Hand-tracking studies have pointed to two dissociable CSE processes: motoric inhibition, reflected in movement initiation times, and controlled response selection, reflected in movement times and reach trajectories (Erb & Marcovitch, 2018, 2019; Erb et al., 2016). Developmental differences further underscore this dissociation, with movement initiation times reaching adult-like levels by ages 10-12, whereas movement times and reach trajectories continue to improve into early adulthood (Erb & Marcovitch, 2019).

fMRI studies of CSE have highlighted two key brain areas: the anterior cingulate cortex (ACC), for conflict detection, and the dorsolateral prefrontal cortex, for control adjustments that enhance subsequent task performance (Carter & van Veen, 2007; Kerns et al., 2004). Egner and Hirsch (2005) also found amplified task-relevant visual cortex activation following conflict, suggesting attentional filtering mechanisms. EEG studies complement this work by implicating mid-frontal theta-band oscillations (∼4–8 Hz) as neural markers of conflict processing, predicting subsequent performance adjustments (Cavanagh & Frank, 2014; Cohen & Donner, 2013; Pastötter et al., 2013). Distinct theta oscillations, including transient, stimulus-locked and sustained, response-locked activity, reinforce the complexity of underlying processes (Töllner et al., 2017). Additionally, N2 and P3 ERP components have shown reduced amplitudes following conflict, indicative of maintained heightened control states (Clayson & Larson, 2011; Larson et al., 2012).

More recently, EEG decoding methods have enabled more sensitive analyses of perceptual (Nemrodov et al., 2018; Shoura et al., 2025) and cognitive phenomena (Bae & Luck, 2018; Fahrenfort et al., 2018) beyond traditional ERP approaches. Using such methods, Lee et al. (2025) have found selective distractor suppression following conflict, particularly in the beta band, consistent with theories associating beta activity with cognitive set maintenance (Engel & Fries, 2010; Wiesman et al., 2020).

Here, we combine EEG decoding techniques with a release-and-press version of the flanker task to probe the CSE both behaviorally and neurally. Using high-density EEG we tested whether neural patterns sensitive to trial congruency were modulated by prior congruency, essentially assessing neural carryover corresponding to the CSE. Time-resolved decoding and cross-temporal generalization tracked the temporal stability of these patterns, clarifying when prior-trial influences emerge. Finally, scalp-based searchlight decoding identified topographic regions associated with sequence effects.

This multipronged approach clarifies when and under what conditions neural CSEs emerge, how they depend on response repetition, and whether they reflect transient or sustained representations. By integrating cognitive control tasks with EEG decoding, we examine the spatio-temporal dynamics of the CSE and test its correspondence with behavioral performance in a two-response version of the arrow flanker task.

## Methods

### Participants

A total of 30 young adults (age range: 18-25 years; 23 females) from the University of Auckland community volunteered for the study. All participants were right-handed, had normal or corrected-to-normal vision, and no diagnosis of cognitive impairments. Six additional participants were excluded due to unmet eligibility, task performance errors, or excessive EEG noise from eye or body movements. Participants were compensated with either course credit or a $25 supermarket voucher. All protocols were approved by the University of Auckland Human Participants Ethics Committee (UAHPEC).

A power analysis (G*Power 3.1; Faul et al., 2009) for an effect size of 𝜂*_p_*^2^= 0.28 (see Erb & Marcovitch, 2018) with 95% power and 5% Type I error rate indicated that a sample size of 29 participants is needed, rendering our sample size adequate.

### Experimental procedures

Participants completed the task in a dimly lit, electrically shielded booth. Task stimuli were presented on a centrally positioned monitor approximately 57 cm from the participant. The monitor was configured to a 640 × 480-pixel resolution with 60Hz refresh rate. Participants responded using a response box (36 cm × 23 cm) placed centrally on the table, equipped with two response buttons and a central “start” button. The E-Prime® software (version 2.0.8.74, Psychology Software Tools; Schneider et al., 2002) managed synchronization of behavioral and EEG recordings, stimulus presentation, and data collection.

The experimental procedure closely followed that described by Smith et al. (2022). Participants performed a computerized two-alternative forced-choice version of the Eriksen flanker task with arrow stimuli. Each trial began with participants pressing and holding the center button on a response box. After 1000ms, a central fixation cross appeared for a randomized period between 800ms and 1200ms. Following the fixation, an array of five evenly spaced arrows (2.2 cm × 2.2 cm each) appeared, with participants instructed to respond to the central arrow while ignoring the flanking distractors. Responses were made by releasing the center “start” button and pressing one of the two response buttons, aiming for both speed and accuracy – see Fig. 1.

**Figure 1.**
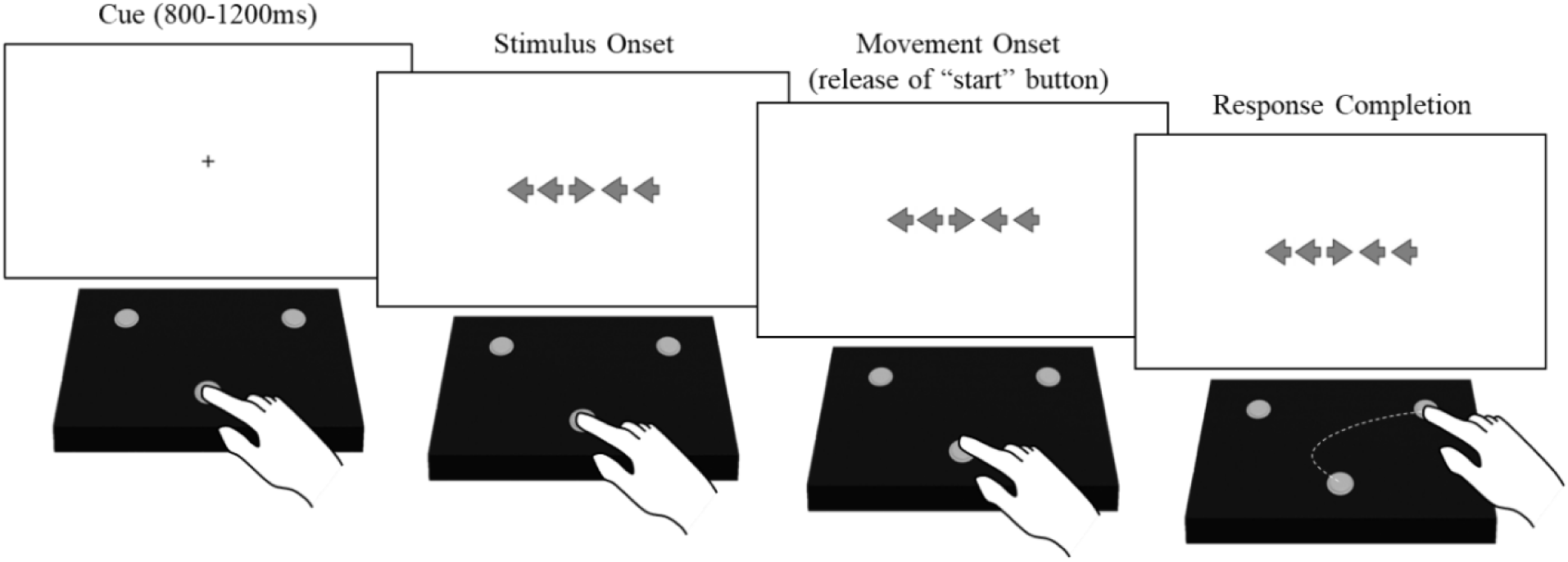
Illustration of task procedure. Participants begin each trial by pressing and holding a “start” button located at the bottom center of a response box. After the stimulus is presented, they release the “start” button and respond by pressing one of two response buttons positioned at the top of the box. Initiation time (IT) is defined as the interval between stimulus onset and the release of the “start” button, while movement time (MT) is the interval between releasing the “start” button and pressing a response button (reproduced from Smith et al., 2022 with permission).

Participants first read written instructions, received a verbal explanation, and watched a demonstration by the experimenter using the response box on the initial trial of a practice block. Then, they completed 16 practice trials, with feedback provided as needed (e.g., “Hold the start button until the arrows appear”). If the center button was released prematurely during fixation or if a response was made too early (within 1000 ms), an on-screen message (“Too early lift”) prompted a trial restart. To proceed to the main experiment, participants needed to correctly complete at least 75% of practice trials.

The main task consisted of 8 blocks of 76 trials each, totaling 608 trials. After each block, a 10s on-screen countdown was displayed before the next block. Each block included an equal number of congruent and incongruent trials, with an equal split of trials requiring left and right responses.

### EEG acquisition and preprocessing

EEG recordings were conducted using 128-channel Ag/AgCl electrode nets (Electrical Geodesics Inc.). Data were recorded at a 1000Hz sampling rate and downsampled to 250Hz. Electrode impedances were adjusted below 40 kΩ.

Data were pre-processed using MATLAB (2021a, The MathWorks Inc.) and the EEGLab toolbox (Delorme & Makeig, 2004). To reduce slow signal drift, data were initially high-pass filtered at 0.1Hz using a bi-directional Butterworth filter. Noisy electrodes were identified, and their signal was approximated by spherical-spline interpolation. Then, data were re-referenced to a common average.

Adaptive mixture independent component analysis (AMICA; Palmer et al., 2011) was applied to the data. To mitigate aliasing from high-frequency noise, a low-pass filter at 30 Hz was subsequently applied using a standard FIR filter. The gamma frequency band (30–100 Hz), associated with bottom–up processing (Buschman & Miller, 2007) and motor execution kinematics (Tatti et al., 2022), was excluded from analysis. Artifacts were automatically detected using ICLabel (Pion-Tonachini et al., 2019) and then manually inspected to confirm appropriate component removal. The data were segmented into epochs for each trial, spanning -1000ms to 3000ms after stimulus onset, a duration chosen to capture both movement initiation and response completion, even in trials with extended initiation or movement times.

After linear detrending of each epoch, the dataset was duplicated into stimulus-locked (200ms before to 999ms after stimulus onset) and initiation-locked (1000ms before to 999ms after movement initiation) versions. Each electrode in these epochs underwent baseline correction, using a -200 to 0 pre-stimulus period. Lastly, epochs were screened for extreme voltages, rejecting epochs with any channel voltage exceeding ±150μV at any time point.

## Statistical Analyses

### Behavioral data

Consistent with prior studies, we defined initiation time (IT) as the interval between stimulus onset and the start of movement (marked by release of the start button), movement time (MT) as the interval between movement initiation and the response (marked by pressing the response button), and total time (TT) as the total time from stimulus onset to response completion. A three-way ANOVA (previous congruency: congruent, incongruent × current congruency: congruent, incongruent × response type: switched, repeated) was performed separately for each reaction time (RT) metric. Key markers, including movement initiation (release of the “start” button) and response completion (press of the response button) were synchronized with the EEG recordings. Finally, and to directly probe the CSE, we compared the difference in TT as well as MT between previously congruent and previously incongruent trials, for both repeated and switched responses, via paired t-tests.

The first trial of each block was excluded from analysis as well as trials containing errors. To control for post-error adjustments (Danielmeier & Ullsperger, 2011), trials immediately following an error were also excluded. Further, trials with response times exceeding 2000ms and initiation times below 100ms were removed to account for potential attentional lapses or off-task interruptions (e.g., posture adjustments). Subsequently, trials exceeding ±3SDs from each participant’s TT mean were also excluded. Overall, 13.21% of trials were excluded from data analysis.

To strengthen confidence in significant effects and clarify nonsignificant ones, we used Bayesian hypothesis testing (JASP). BF₁₀ values were interpreted in favor of the alternative (1–3 = anecdotal, 3–10 = moderate, 10–30 = strong, >30 = very strong) or the null (1/3–1 = anecdotal, 1/10–1/3 = moderate, 1/30–1/10 = strong, <1/30 = very strong) (Schonbrodt & Wagenmakers, 2018). Analyses relied on JASP’s default priors (e.g., uniform for ANOVAs, Cauchy for t-tests).

### Temporally cumulative analysis

Data was further processed and resampled for pattern analysis purposes. For each condition within each block, we randomly selected two trials, averaging them to construct a more robust pseudotrial. This procedure was repeated ten times, generating 80 pseudotrials per block. Specifically, this yielded 10 averaged observations across eight conditions: cC-r, cC-s, cI-r, cI-s, iC-r, iC-s, iI-r, iI-s, where i/c refers to previous trial in/congruency, I/C to current trial in/congruency and r/s to repeated/switched responses.

To construct spatiotemporal patterns, time points beginning at 50 ms post-stimulus onset were concatenated with data from all 128 channels, resulting in 30,464 features per observation (128 channels × 238 time points). This yielded 10 patterns for each condition, and each of the eight blocks. Temporally cumulative analysis (TCA) relies on spatiotemporal information in such patterns to boost decoding accuracy at the cost of temporal specificity (Nemrodov et al., 2018) – but see complementary time-resolved analyses below.

We performed pairwise classification using a linear SVM (c = 1) with leave-one-block-out cross-validation. The resampling procedure described above along with classification was repeated 100 times to ensure stable accuracy estimates, with the final classification accuracy calculated as the average across iterations. Chance-level accuracy was determined via one-sample Wilcoxon signed-rank test against chance (50%).

A one-way ANOVA (3 conditions: previous congruency, current congruency, response type) was performed on the decoding accuracies. Additionally, a 2 × 2 ANOVA (response type: switched, repeated × previous congruency: congruent, incongruent) was performed on the decoding accuracies of current trials to investigate the CSE. For completeness, this process was also repeated for the decoding of previous congruency and response type.

### Neural dynamics

To examine decoding accuracy over the duration of a trial, time-resolved analysis (TRA) was conducted using overlapping five-timepoint sliding windows (∼10ms), with a stride of one timepoint. Significance was assessed with the aid of cluster-based tests (1000 permutations; cluster-forming threshold, *p* < .05). Analyses were conducted separately for stimulus-locked and initiation-locked data to capture distinctions in neural dynamics linked to stimulus presentation and response initiation. This analysis was then repeated for the decoding accuracies of congruent trials as a function of previous congruency and response type to investigate the CSE.

To assess the stability of neural patterns across time, cross-temporal generalization was performed on current congruency decoding as a function of previous congruency and response type. For this analysis, training was conducted on data from a specific temporal window, and testing was conducted on all other temporal windows. This generated a cross-temporal decoding matrix, where each cell represents classification accuracy for a particular combination of training and testing windows. Significance was assessed using Wilcoxon signed-rank tests (FDR-corrected across time points, *q* < .05).

### Frequency-band analysis

We examined congruency decoding within specific EEG frequency bands: theta (4–8 Hz), alpha (8–12 Hz), low beta (12–20 Hz), and high beta (20–30 Hz). The beta band was separated into low and high to explore their distinct roles in cognitive control and congruency sequence effects (Lee et al., 2025). Because the optimal time window for such an analysis was unclear, we used TRA results (see Neural dynamics) to pinpoint relevant intervals. A 2 × 4 ANOVA (previous congruency × four frequency bands) was performed on the decoding accuracies of current congruency for repeated trials. To test the importance of theta band for CSE (Cavanagh & Frank, 2014; Cohen & Donner, 2013; Pastötter et al., 2013) we directly assessed the advantage of current congruency decoding when preceded by previously congruent over incongruent trials (one-tailed Wilcoxon signed-ranked test).

### Searchlight analysis

To assess differences in decoding accuracy across scalp regions, classification was conducted at the location of each electrode, including its five nearest neighboring electrodes. Thus, this approach uses small clusters of electrodes to estimate decoding accuracy in spatially restricted regions. Separate analyses were conducted for switched and repeated responses across two key time windows: 450-500ms and 500-550ms. Significance was assessed using Wilcoxon signed-rank tests (FDR-corrected across time points, *q* < .05).

### Brain-behavior correlations

Finally, to examine the relationship between neural and behavioral measures of conflict processing, we computed Pearson correlations between current congruency decoding accuracy and behavioral congruency effects (i.e., RT differences between incongruent and congruent trials) for TT, IT, and MT.

To examine whether neural markers of CSE corresponded with behavioral CSE effects, we computed, for each participant, the difference in current congruency decoding between previously congruent and incongruent trials, separately for repeated and switched responses. These neural CSE scores were then correlated with behavioral CSE scores computed analogously for TT, IT, and MT.

## Results

### Behavioral performance

TTs were analyzed to assess differences in overall task performance across conditions (Fig 2A). The analysis revealed an effect of current congruency (*F*(1,29) = 203.662, *p* < .001, 𝜂*_p_*^2^= .875). Consistent with a CSE, the interaction between previous and current congruency (*F*(1,29) = 5.518, *p* = .026, 𝜂*_p_*^2^= .16) was significant. Further interactions between previous congruency and repetition type (*F*(1,29) = 9.816, *p* =.004, 𝜂*_p_*^2^= .253), and between the three variables (*F*(1,29) = 13.462, *p* < .001, 𝜂*_p_*^2^= .317) were also significant, with Bayesian analysis indicating that the best model includes all main effects and interactions (*BF*_10_ = 1.91 × 10^14^). Accordingly, to validate the CSE, we directly compared the size of the current congruency effect after congruent versus incongruent trials, separately for switched and repeated trials. A robust CSE – manifested as a larger current congruency effect after congruent trials – was observed only on repeated trials (*t*(29) = 4.203, *p* < .001, *d* = .767, *BF*_10_ = 122.733), but not on switched trials (*t*(29) = 0.362, *p* = .72, *d* = .066, *BF*_10_ = 0.207).

**Figure 2.**
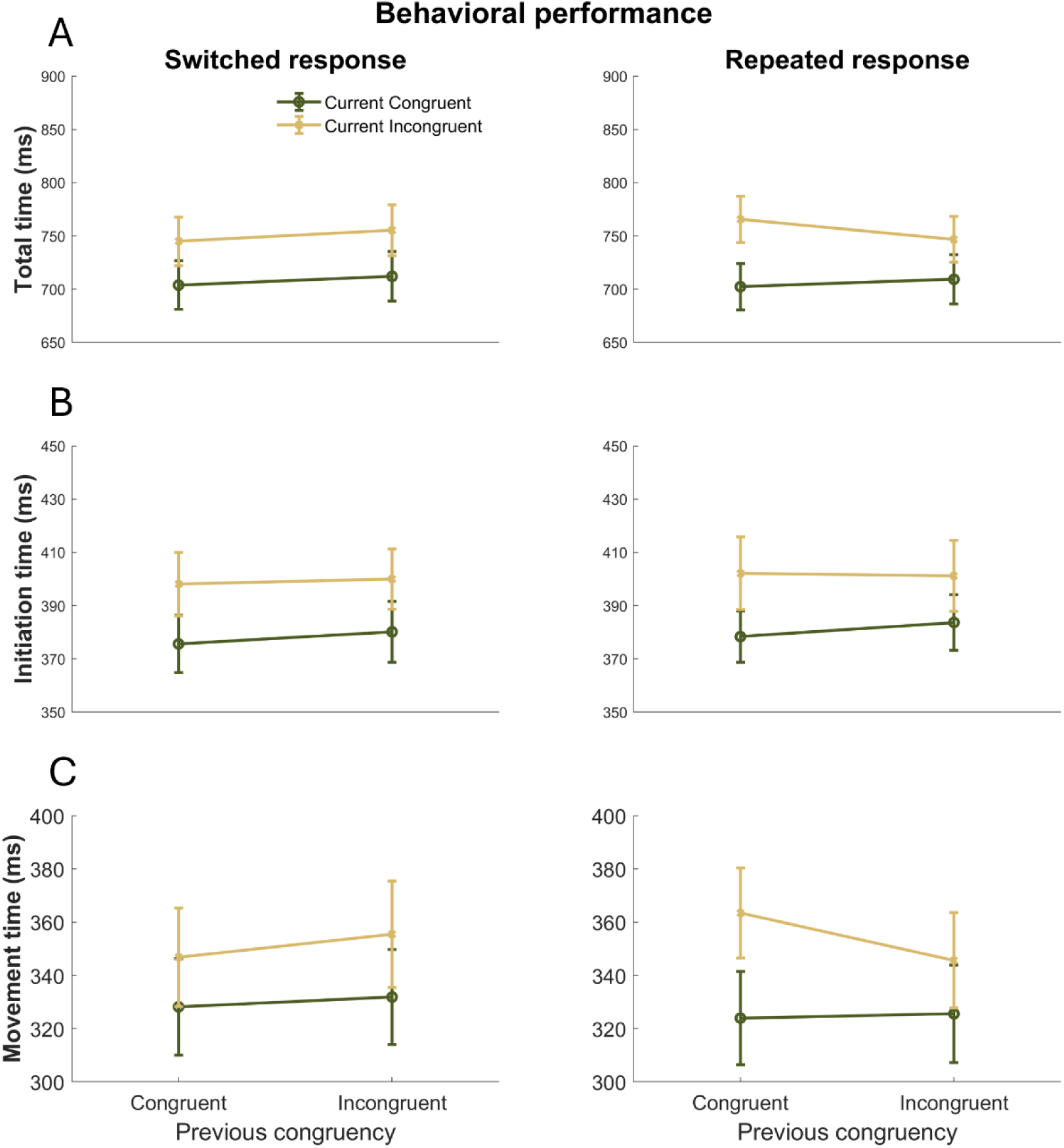
(A) Total time, (B) initiation time, and (C) movement time in the two-response flanker task. Facilitation for iI relative to cI trials (incongruent trials preceded by an incongruent trial vs. a congruent trial) was found for total time and movement time of repeated responses (error bars show ±1SE).

ITs were examined to explore differences in the initial response phase between conditions (Fig. 2B). There was a main effect of both previous (*F*(1,29) = 10.214, *p* = .003, 𝜂*_p_*^2^ = .26) and current (*F*(1,29) = 48.631, *p* < .001, 𝜂*_p_*^2^= .626) congruency but no significant interactions. Bayesian analysis revealed that the best model includes only previous and current congruency effects (*BF*_10_ = 174,658.67).

MTs were analyzed to evaluate differences in the execution phase of the response across conditions (Fig. 2C). Similar to TT results, the effect of current congruency (*F*(1,29) = 79.75, *p* < .001, 𝜂*_p_*^2^= .733) as well as the interaction between previous and current congruency (*F*(1,29) = 4.867, *p* = .035, 𝜂*_p_*^2^= .144) were significant. Further interactions between previous congruency and repetition type (*F*(1,29) = 11.321, *p* =.002, 𝜂*_p_*^2^= .281), and between the three variables (*F*(1,29) = 9.875, *p* = .004, 𝜂*_p_*^2^ = .254) were also significant, with Bayesian analysis indicating that the best model includes all main effects and interactions (*BF*_10_ = 1.30 × 10^9^). Importantly, a robust CSE was observed only on repeated trials (*t*(29) = 3.581, *p* = .001, *d* = .654, *BF*_10_ = 27.594), but not on switched trials (*t*(29) = 1.058, *p* = .299, *d* = .193, *BF*_10_ = .324), consistent with its modulation by stimulus-response (S-R) binding during movement (Erb & Marcovitch, 2018).

Given that accuracy was at ceiling (M = 99.16%, SD = 0.01%), no additional analyses were conducted on error rates.

### Temporally cumulative decoding

Decoding accuracies for current congruency (*W* = 465, *p* < .001, *BF*_10_ = 3.50 × 10^11^) and response type (*W* = 463, *p* < .001, *BF*_10_ = 3.96 × 10^6^) but not for previous congruency (*W* = 290, *p* = .245, *BF*_10_ = .546) were significantly above chance. The three decoding estimates were significantly different from each other (*F*(2,58) = 88.222, *p* < .001, 𝜂*_p_*^2^= .753, *BF*_10_ = 1.347 × 10^18^). All pairwise comparisons were significant (all *p’*s < .001, all *d*s ≥ 1.248, all *BF*_10_ values ≥ 6451.767) showing a clear advantage for decoding current congruency over the other two types of decoding as well as an advantage for response type over previous congruency – see Fig. 3A.

**Figure 3.**
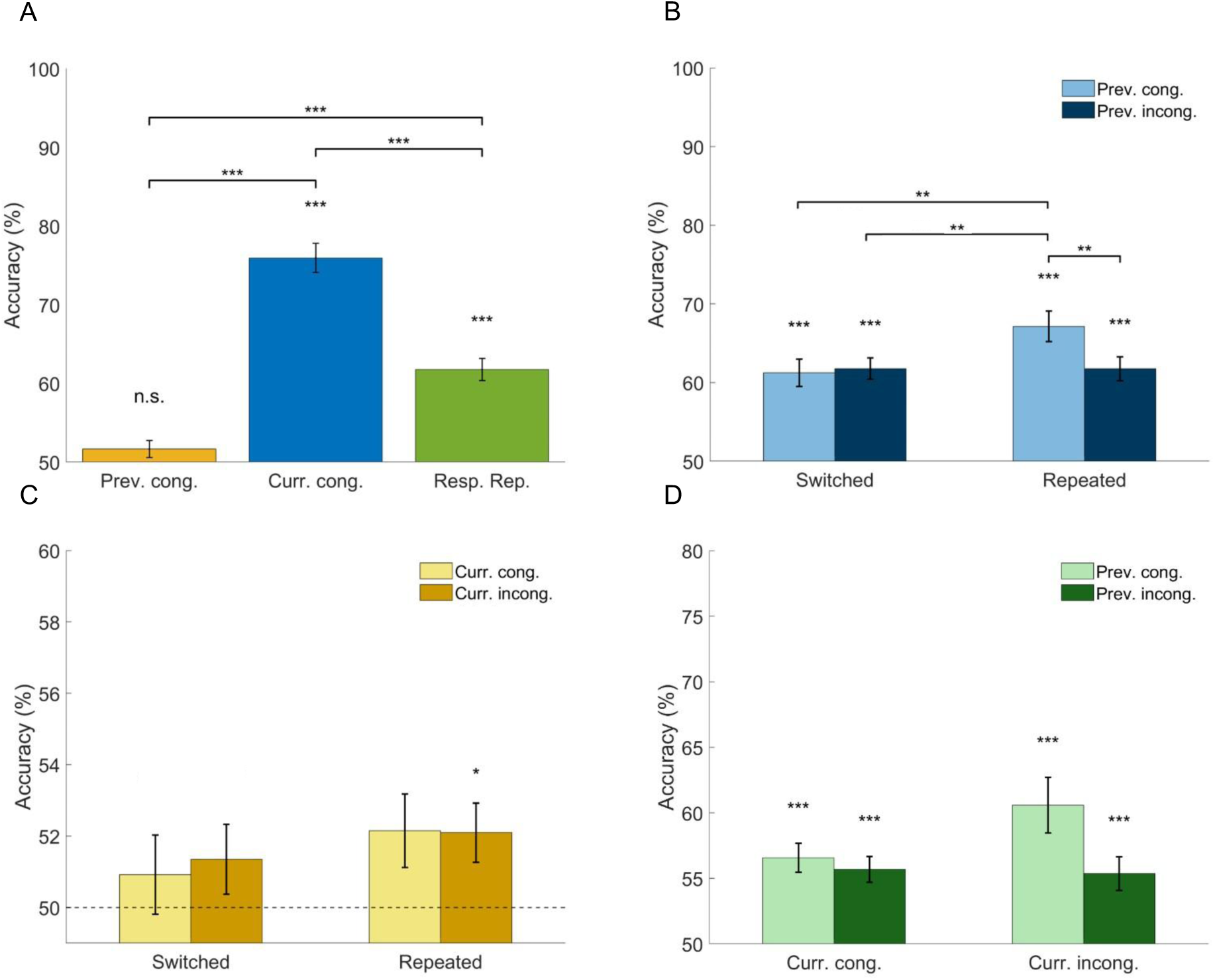
(A) Temporally cumulative decoding for previous congruency, current congruency, and response type showed reliable decoding of current congruency and response type. (B) Decoding of current congruency yields higher accuracy for repeated responses following previously congruent trials relative to other conditions. (C) Decoding of previous congruency yields significantly above-chance decoding only for repeated, currently incongruent trials. (D) Decoding of response type yields significantly above-chance decoding for all conditions (\*\*\**p* < .001, \*\**p* < .01, \**p* <. 05).

Next, current congruency decoding was analyzed across previous congruency (congruent, incongruent) and response type (switched, repeated) via repeated measures ANOVA – see Fig. 3B. The analysis revealed a significant main effect of previous congruency (*F*(1,29) = 5.683, *p* = .024, 𝜂*_p_*^2^ = .164) and response type (*F*(1,29) = 5.259, *p* = .029, 𝜂*_p_*^2^ = .154), as well as an interaction (*F*(1,29) = 9.123, *p* = .005, 𝜂*_p_*^2^= .239). Bayesian analysis revealed that the best model includes all main effects and interaction (*BF*_10_ = 56.19). Notably, a previous congruency advantage was found for repeated responses (cC-r vs. cI-r; *t*(29) = 3.812, *p* = .002, *d* = .595) but not for switched responses (cC-s vs. cI-s; *t*(29) = -.379, *p* > .999, *d* = -.059). This indicates a neural CSE only for repeated responses, mirroring the behavioral results above.

Next, we decoded previous congruency across levels of current congruency and response type – see Fig. 3C. No main effects or interaction were significant. The best Bayesian model included response type only (*BF*_10_ = .374). We found above-chance decoding for incongruent repeated trials (cI-r vs. iI-r; W = 341, p = .026, *BF*_10_ = 2.817) but not for the other decoding conditions. This suggests that information about prior congruency may persist during a current trial but is modulated by current congruency and response type.

For completeness, we also decoded response type as a function of current and previous congruency – see Fig. 3D. This revealed a significant advantage for previously congruent over incongruent trials (*F*(1,29) = 5.726, *p* = .023, 𝜂*_p_*^2^= .056) as well as a significant interaction between current and previous congruency (*F*(1,29) = 4.208, *p* = .049, 𝜂*_p_*^2^= .028). However, no post hoc tests yielded significant results. Bayesian analysis revealed that the best model includes previous congruency only (*BF*_10_ = 1.905).

While TCA provided robust decoding by aggregating EEG signals across a large interval, it sacrifices temporal specificity regarding when congruency effects emerge. To address this, we performed TRA decoding, aiming to characterize the temporal dynamics of congruency-related neural activity.

### Neural dynamics

To assess the temporal profile of congruency effects, decoding was performed using successive ∼10-ms intervals rather than a single large time window spanning the trial – see Fig. 4A. For stimulus-locked signals, significant decoding of current congruency was present between ∼200-500ms, with a peak around 450ms, and also between ∼ 600-900ms. Decoding of response type was less robust, but was significant between ∼300-750ms, with a peak also around 450ms. In contrast, no significant time windows were observed for previous congruency.

**Figure 4.**
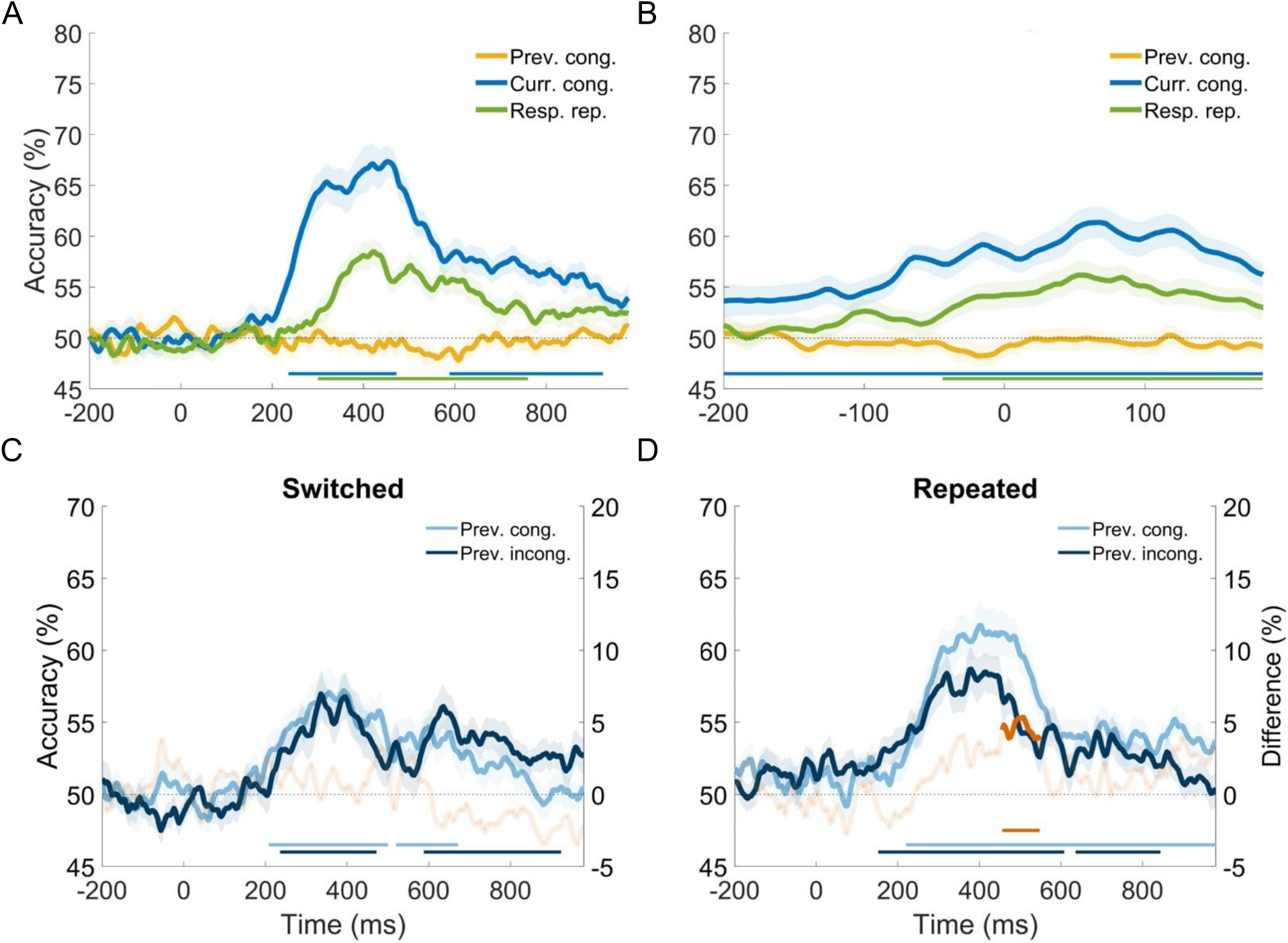
(A) Time-resolved decoding for stimulus-locked trials revealed intervals of above-chance decoding for current congruency and response type but not for previous congruency. (B) Decoding for movement initiation-locked trials yielded significant but less robust findings. (C) Current congruency decoding for switched trials found no significant differences between the previously congruent and incongruent trails. (D) Current congruency decoding for repeated trials revealed a significant advantage for previously congruent trials between ∼450-550ms (cluster test, *p* < .05).

The same analysis was repeated with signals locked to movement initiation - see Fig. 4B. Decoding of current congruency was significantly above chance from 200ms pre-initiation to ∼200ms after initiation, with a peak around 50ms. Response type decoding was significant around -50ms, tapering off around 200ms. Consistent with the stimulus-locked analysis, no significant time windows were observed for previous congruency.

Next, we assessed current congruency decoding separately for switched and repeated responses – see Fig. 4C, D. Switched responses for both previously congruent (cI-s vs. cC-s) and incongruent trials (iI-s vs. iC-s) yielded multiple significant windows but no difference between the two. In contrast, repeated responses yielded a significant advantage for previously congruent over incongruent trials between ∼450ms to 550ms – see Fig 4D.

Thus, TRA identified a specific interval (∼450–550ms) associated with the CSE. Beyond its theoretical significance, this result also provides an empirical basis for selecting a targeted window for subsequent frequency-band analyses.

### Frequency band analysis

Guided by the temporal profile of CSE on repeated trials (Fig. 4D), we computed a more targeted TCA for the two conditions, separately for four frequency bands: theta, alpha, low beta, and high beta. We considered as a window of interest the interval between 200-600ms, during which we noticed a numeric advantage for previous congruency (Fig. 4D). We note that this larger window, aiming to capture reliably lower-frequency information, includes the significant difference between ∼450-550ms found above.

Decoding revealed above-chance accuracies in the theta and alpha bands (all *W*s ≥ 424, all *p*s < .001, all *BF*_10_ values ≥ 97.567) but not in the beta bands (all *ps* ≥ .368) – see Fig 5. The hypothesis that the decoding advantage for previously congruent trials stems specifically from theta activity found support in our data (*W*(29) = 338, *p* = .04, *d* = .363, *BF*_10_ = 3.442, uncorrected). No such advantage was observed in the alpha, low beta, or high beta bands (all *ps* ≥ .231, uncorrected).

**Figure 5.**
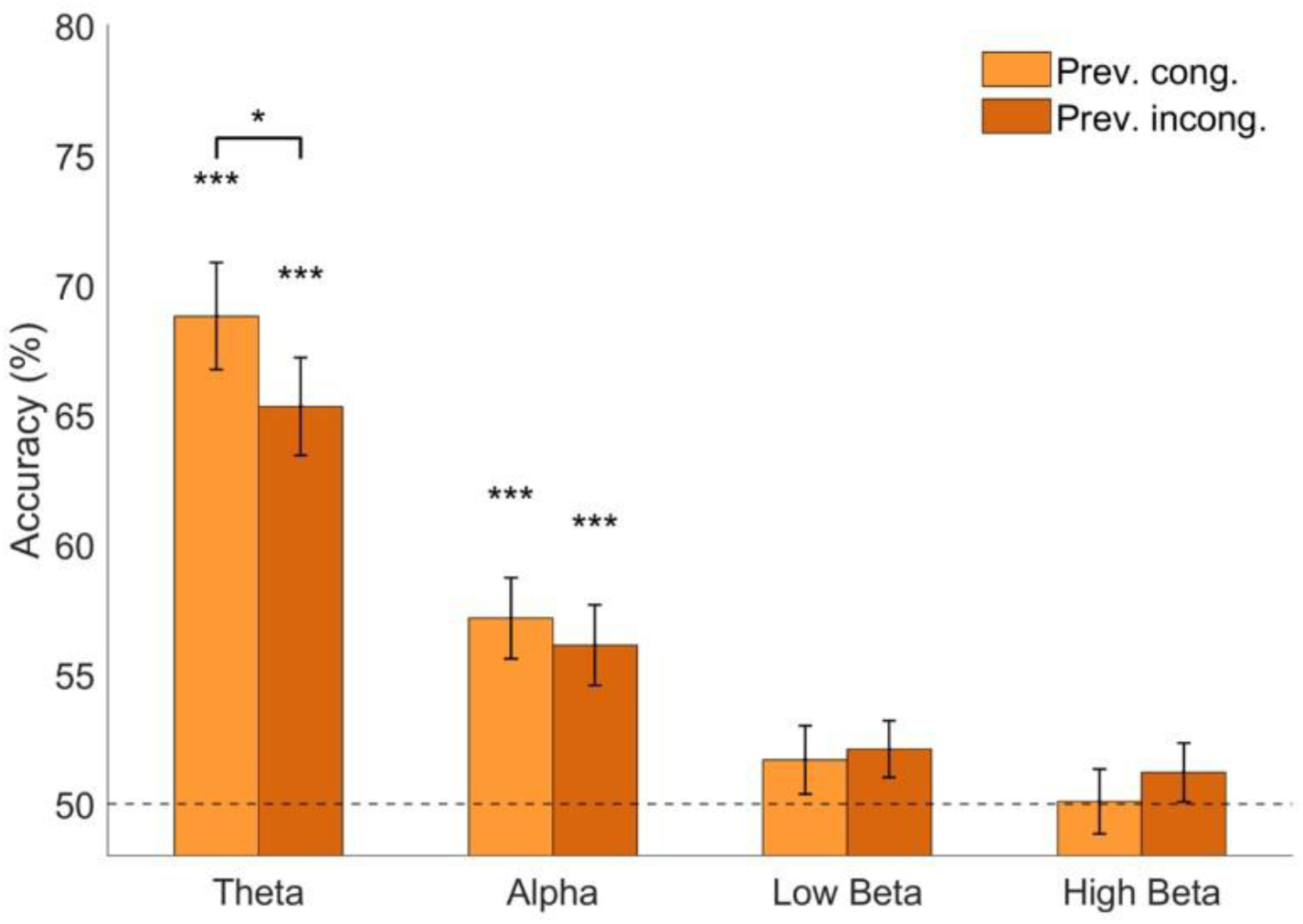
Temporally cumulative decoding of current congruency for repeated trials across four different frequency bands. Higher decoding for previously congruent over incongruent trial was found in theta, but not in other frequency bands (\*\*\**p* < .001; \**p* < .05 uncorrected).

### Cross-temporal generalization

An analysis of current congruency decoding across time, split by previous congruency and response type, revealed limited cross-temporal generalization (FDR-corrected, *q* < .05; Fig. 6). Significant decoding was restricted to a narrow region along the diagonal for previously congruent repeated trials (Fig. 6B), with a weaker but similar pattern for previously incongruent repeated trials (Fig. 6D).

**Figure 6.**
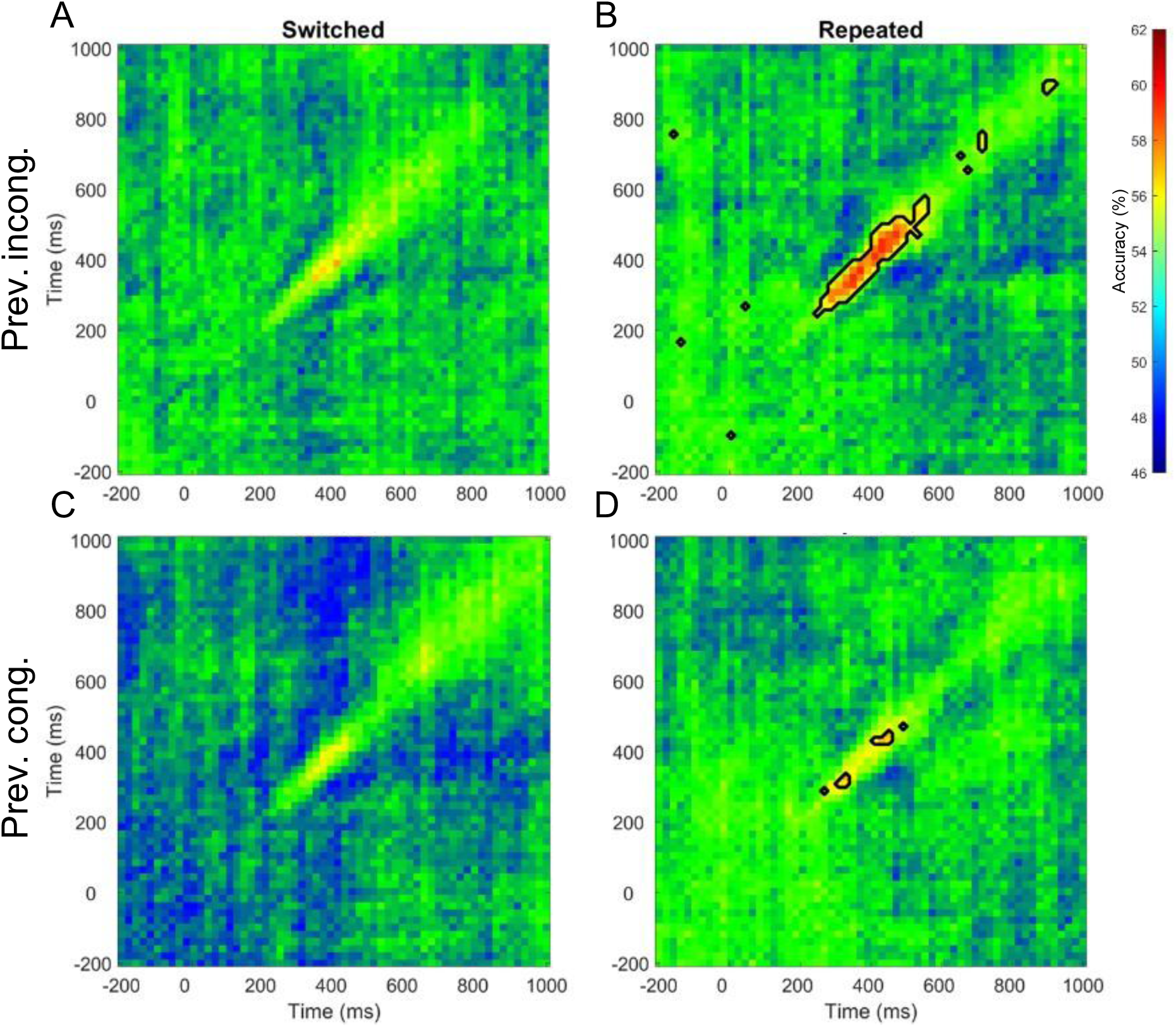
Cross-temporal generalizability analysis shows limited generalization for repeated responses near the diagonal and no significant generalization for switched responses. The horizontal axis corresponds to testing timepoints, and vertical axis corresponds to training timepoints (FDR corrected across timepoints, *q* < .05).

This pattern supports a chain-of-generators architecture (King & Dehaene, 2014), with no evidence of pattern reactivation. Instead, the results suggest transient, sequential neural representations. This is consistent with control being implemented via a cascade of evolving stages rather than sustained or recurrent top-down feedback (Hinault et al., 2019).

### Searchlight decoding

To characterize the spatial profile of decoding across scalp regions, we performed searchlight decoding of current congruency for repeated responses across the significant window identified by TRA. Further, we aimed to pinpoint the regions responsible for the current congruency advantage of previous congruency over incongruency.

To examine if and how the advantage above emerges over time, we divided the interval of interest in two halves: 450–500ms and 500–550ms. During the first window, decoding of current congruency following previously incongruent trials exhibited significant decoding across all regions except for medial occipital regions, while the second time-window showed less robust decoding, spanning central, frontal, and occipital regions (Wilcoxon signed-rank test, FDR-corrected, *q* <.05). Decoding current congruency following previously congruent trials revealed significant decoding throughout all regions for both time windows, with more robust decoding for the 450-500ms time-window – see Fig 7A, B. For the first window, electrodes corresponding to Fp1 and Fp2 yielded the highest decoding accuracy.

**Figure 7.**
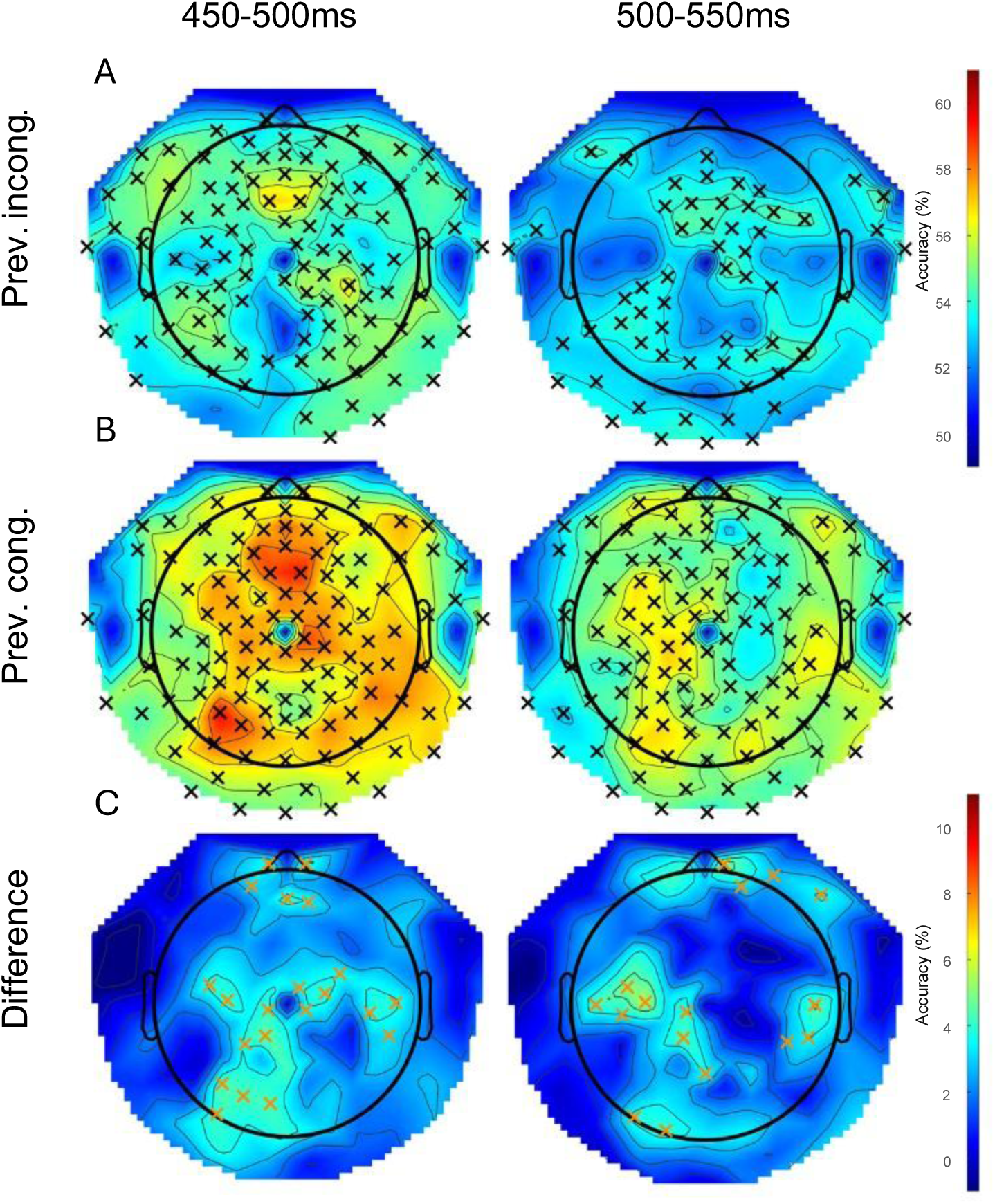
Searchlight analysis of current congruency for repeated responses during two time windows when previous trials were (A) incongruent, and (B) congruent (*q* < 0.05, FDR-corrected). (C) The difference between the two searchlight maps points to frontal, central and left occipital regions (Wilcoxon signed-rank test, *p* < 0.05, uncorrected).

The difference between previously congruent and incongruent searchlight maps, pointed to right frontal, central, and left occipital regions for both time windows (Wilcoxon signed rank test, *p* < .05, uncorrected) – see Fig 7C. However, no regions reached significance after multiple-comparison correction.

To address potential confounds of motor responses on the decoding results above, searchlight analysis was repeated for left versus right movements. The decoding patterns observed during both time windows (450-500ms and 500-550ms) differ markedly from those associated with congruency decoding – see supplementary Fig. 1. This suggests that the observed decoding results are unlikely to be driven primarily by motor responses.

### Brain-behavior correlations

We examined whether neural decoding of current congruency (i.e., the congruency effect) was associated with behavioral congruency effects for TT, IT, and MT. No significant correlations were found (all *p*s ≥ .43).

Next, we tested whether neural decoding of current congruency was related to the behavioral CSE. For each participant’s current congruency decoding results, we calculated the average difference between previously congruent and incongruent trials. This was computed for each participant separately for repeated and switched responses. We also quantified the CSE behaviorally for TT, IT, and MT. No significant correlations were found (all *p*s > .22). This lack of association may reflect, at least in part, limited statistical power: our sample size (n = 30) falls well below the recommended threshold (n = 84) for detecting medium-sized effects (*r* = .3; Brysbaert, 2019). Clarifying the link between EEG and behavioral indices of the CSE will require larger samples in future research.

## Discussion

The present findings offer novel insights into the neural basis of the flanker CSE, first reported in the seminal study of Gratton et al. (1992), in particular with regard to the interaction between cognitive control and S-R binding. Our results point to several key observations concerning the neural mechanisms, temporal dynamics, and representational content underlying CSE.

First, behavioral analyses confirm a robust CSE selectively modulated by response type. Congruency effects were significantly diminished following incongruent trials compared to congruent ones, but notably, this reduction only occurred when responses repeated across trials. This pattern aligns with episodic memory accounts of the CSE (Hommel et al., 2004; Mayr et al., 2003), suggesting that the effect is driven by performance costs from an S-R binding conflict (see also Hazeltine et al., 2011 for set-level influences). Importantly, this dependence on response repetition indicates that adaptive behavioral adjustments in the CSE arise not solely from abstract cognitive control processes, but also from automatic episodic retrieval of prior S-R episodes, with binding conflict impairing performance and repetition priming facilitating it (Hommel, 1998; Mayr et al., 2003; Dignath et al., 2019).

Second, our EEG decoding results complement our behavioral findings by revealing neural markers of the CSE that similarly depend on response type. Temporally cumulative analyses demonstrated significantly higher decoding accuracy for current congruency following previously congruent trials compared to incongruent ones, but again only for repeated responses. This congruency-dependent advantage parallels behavioral performance, providing a neural counterpart of the CSE and emphasizing the role of response-bound episodic representations in cognitive control adjustments (Hommel, 2004; Schmidt & Weissman, 2014).

Third, the temporal profile of neural decoding reveals distinct intervals of congruency representation, pinpointing a critical window around 450–550ms post-stimulus onset for repeated responses. This finding aligns with previous work demonstrating mid-latency neural responses associated with attentional filtering and stimulus evaluation in conflict tasks (Egner & Hirsch, 2005; Larson et al., 2012). Additionally, our cross-temporal generalization analysis found limited evidence of sustained or recurrent neural activation patterns, instead suggesting sequential and transient neural representations consistent with cascading stages of cognitive control processing (Hinault et al., 2019).

Fourth, frequency-band analyses underscore the specific contribution of theta-band activity (4–8 Hz) to the neural representation of congruency sequence effects. Theta-band current trial decoding showed a significant advantage for previously congruent repeated responses, corroborating existing EEG studies highlighting frontal theta oscillations as critical markers of cognitive control processes, conflict detection, and adaptive behavior (Cavanagh & Frank, 2014; Cohen & Donner, 2013; Pastötter et al., 2013). Thus, our results reinforce the importance of frontal theta activity in mediating conflict-driven control adjustments, particularly those linked to episodic retrieval processes.

Finally, searchlight decoding pointed to congruency effects over frontal, central, and occipital scalp regions. The broad spatial distribution of these effects, particularly in frontal and central areas during the 450–550ms interval, aligns with neuroimaging evidence implicating ACC and dorsolateral prefrontal cortex in conflict monitoring and subsequent control adjustments (Carter & van Veen, 2007; Kerns et al., 2004). Although specific regions did not survive multiple-comparison corrections, our pattern of results suggests distributed neural networks underpinning CSE, warranting further investigation into precise regional contributions via source localization methods.

Interestingly, despite clear behavioral and neural congruency sequence effects, our analyses did not reveal significant brain-behavior correlations. This dissociation might suggest that neural markers indexed by decoding methods capture cognitive control processes or representations not directly reflected in overt reaction times. Alternatively, this finding might reflect limited variability or ceiling effects in our behavioral data, highlighting the need for future studies employing more sensitive behavioral metrics or varied task difficulty levels to further probe brain-behavior relationships. Finally, this finding may simply be due to our sample size.

From a methodological perspective, this study showcases the advantage of integrating multiple types of EEG decoding techniques to elucidate neural mechanisms underlying cognitive phenomena. TCA allowed robust decoding accuracy by aggregating extensive temporal data, albeit sacrificing temporal specificity, whereas complementary TRA effectively pinpointed critical processing windows. Future research could benefit from similarly integrated methodological approaches, employing complementary decoding methods to capture both the stability and temporal dynamics of neural processes.

A few caveats warrant mention here. First, while our release-and-press response box paradigm provides rich data on the temporal dynamics of decisions, it also inherently couples cognitive conflict with motor execution. Our analysis of hand movement (i.e., left vs. right decoding) and the lack of significant brain-behavior correlations suggest that the effects reported above are not due to trivial motor artifacts. Further, while aspects of motor kinematics such as velocity and acceleration are intertwined with cognitive control, we note that movement speed has been linked to beta-band activity (Zhang et al., 2020), while velocity has been associated with the gamma band (Tatti et al., 2022). In contrast, the CSE was associated with the theta band, as hypothesized based on its relevance for cognitive control (Cavanagh & Frank, 2014; Cohen & Donner, 2013).

Further, our sample consisted of young adults performing at >99% accuracy. It remains to be seen how these dynamics might differ in scenarios with more errors or in populations with differing cognitive control efficacy. Error-related adjustments, for instance, might engage partly different mechanisms than conflict-related adjustments (though we excluded post-error trials here). Investigating the CSE in individuals with executive function impairments could test whether the absence of a neural CSE (e.g., lack of theta upregulation or lack of context-specific pattern changes) predicts poorer adaptation behaviorally.

To conclude, by combining behavioral, neural, and temporal analyses, the current study significantly advances our understanding of congruency sequence effects. Our findings underscore the crucial interplay between memory-based S-R binding and cognitive control mechanisms, revealing dynamic neural representations sensitive to prior trial congruency and response repetition. These insights refine existing theoretical accounts and open avenues for further exploration into the distinct yet interacting cognitive and mnemonic processes shaping adaptive behavior.

---

**Supplementary Figure 1.**
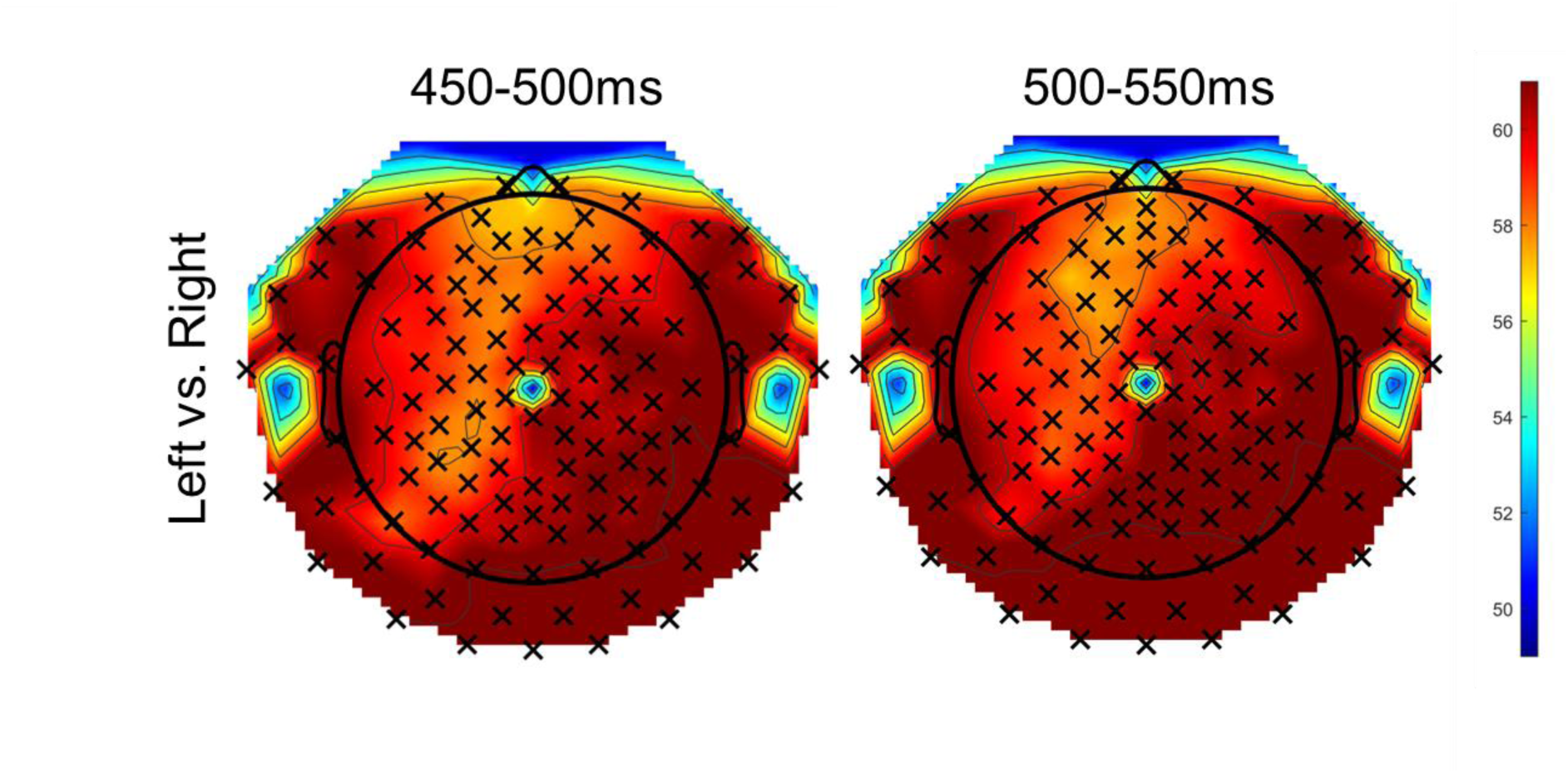
Searchlight analysis of hand movement (*q* < 0.05, FDR-corrected) reveals distinct patterns from that of current congruency.

